# DIA-MS based plasma peptidomic workflow for profiling organ-derived peptides

**DOI:** 10.64898/2026.02.02.703436

**Authors:** Yusei Okuda, Ryo Konno, Tomomi Taguchi, Makoto Itakura, Takashi Matsui, Takeshi Miyatsuka, Osamu Ohara, Yusuke Kawashima, Yoshio Kodera

**Affiliations:** Department of Physics, School of Science, Kitasato University, Kanagawa, Japan; Department of Applied Genomics, Kazusa DNA Research Institute, Chiba, Japan; Department of Diabetes, Endocrinology and Metabolism, Kitasato University School of Medicine, Kanagawa, Japan; Department of Biochemistry, Kitasato University School of Medicine, Kanagawa, Japan; Center for Disease Proteomics, School of Science, Kitasato University, Kanagawa, Japan

**Keywords:** peptidomics, peptidome, DIA-MS, mouse, plasma, organs

## Abstract

Plasma contains diverse bioactive peptides that play crucial roles in maintaining homeostasis and regulating disease responses. However, the presence of peptides derived from high-abundance proteins such as albumin makes comprehensive analysis of native peptides secreted by organs challenging. This study aimed to establish a highly sensitive plasma peptidomic approach by combining data-independent acquisition (DIA) with spectral libraries of plasma and organs. First, peptides were extracted from plasma and eleven organ types using a high-yield peptide extraction method, the differential solubilization method. These peptides were then measured via data-dependent acquisition (DDA) analysis using a timsTOF HT for constructing empirical spectral library. Subsequently, DIA-MS data from plasma samples were measured and analyzed using this spectral library. This strategy achieved identification of, on average, over 5,500 peptides per run, with over 2,000 organ-derived peptides including 19 known bioactive peptides.

The novel strategy proposed here enables highly sensitive quantitative analysis of organ-derived peptides in plasma, linking them to their secreting organs. It is expected to substantially contribute not only to the discovery of biomarkers and novel bioactive peptides but also to elucidating the pathophysiology of systemic diseases.

## Introduction

Plasma contains a wide variety of bioactive peptides, including hormones and neuropeptides, that are secreted from various organs. These molecules play central roles in maintaining homeostasis and modulating disease responses. For instance, well-known peptides like insulin, endorphins, and oxytocin are involved in a broad range of biological phenomena, including metabolic regulation, pain control, and antistress effect, respectively[1], [2], [3]. Consequently, bioactive peptides are gaining increasing attention as diagnostic markers and therapeutic targets and are now widely applied as biomarkers and peptide-based drugs [4], [5]. However, the currently identified bioactive peptides in blood represent only the tip of the iceberg; it is believed that a vast number of peptides with uncharacterized functions still exist in the bloodstream [6]. Therefore, the comprehensive identification of novel bioactive peptides in the blood and the elucidation of their functions are critical research imperatives that promise to deepen our understanding of disease mechanisms and lead to the development of innovative therapies.

A major technical barrier in plasma peptidomics is the overwhelming presence of high-abundance proteins, such as albumin and immunoglobulins, which constitute most of the total protein. In addition, many peptides are bound to these high-abundance proteins and must be dissociated before extraction [7]. To address this issue, we previously developed an analytical workflow that combines a high-efficiency peptide extraction method [8], HPLC fractionation, and DDA-MS. Using this approach, we detected more than 10,000 peptides [9], including 22 known bioactive peptides, from human plasma by performing around 200 DDA-MS runs. Furthermore, through various functional assays, we successfully identified five novel bioactive peptides [10], [11], [12]. However, in single-shot DDA-MS without prefractionation, most of the detected peptides were derived from high-abundance proteins. Thus, to more comprehensively detect bioactive peptides present at picomolar concentrations in plasma, an analytical method with higher sensitivity than DDA-MS is required.

In recent years, DIA-MS has been increasingly adopted in proteomics as an alternative to DDA-MS owing to its superior sensitivity and reproducibility, enabling comparative analysis using label-free quantification [13]. In peptidomics, however, the situation differs from trypsin-digested proteomics, as a much larger peptide search space must be considered. In peptidomic studies focusing on immunopeptides, the search space can be reduced by restricting analyses to peptides predicted to bind HLA and by limiting peptide length based on biological properties [14]. DIA-MS data of immunopeptides can be analyzed using spectral libraries constructed from DDA-MS data or predicted spectral libraries generated solely from protein sequences [15]. In contrast, in peptidomics, which targets native peptides generated by diverse cleavages from precursor proteins within living organisms, it is difficult to reduce the search space based on constraints such as immunopeptidomics. Consequently, analysis using predicted spectral libraries generated solely from protein sequences is impossible. As a result, a commonly used approach is to construct spectral libraries from DDA-MS data of the same sample and use them to analyze DIA-MS data obtained from that sample [16], [17]. Although this strategy enhances quantifiability, the peptides included in the spectral library are limited to those identified at the DDA-MS level, and the intrinsic sensitivity of DIA-MS may not be fully utilized.

In this study, we devised an approach to analyze DIA-MS data obtained by plasma peptidomic analysis using an empirical spectral library constructed from DDA-MS data of plasma and eleven kinds of organs. This approach enabled the identification and quantification of a greater number of plasma peptides, including organ-derived peptides, compared with other methods. Furthermore, this promises to make it possible to infer which organ secreted peptides in plasma, contributing significantly not only to the discovery of novel bioactive peptides but also to elucidating the mechanisms of systemic diseases.

## Materials and Methods

### Animals and sample preparation

C57BL/6 male mice were purchased from CLEA Japan, Inc. (Tokyo, Japan). All procedures involving animals complied with the guidelines of the National Institutes of Health and were approved by the Animal Experimentation and Ethics Committee of Kitasato University School of Medicine. Eleven mouse organs (lung, spleen, adrenal gland, kidney, heart, liver, muscle, pancreas, cerebellum, hippocampus, and cerebral cortex) were dissected, frozen in liquid nitrogen immediately after dissection, and stored at −80°C. The organ sections were immersed in an ice-cold denaturing solution (7 M urea, 2 M thiourea, 20 mM dithiothreitol, cOmplete Mini Protease Inhibitor Cocktail [Roche, Basel, Switzerland]) and weighed. The samples were diluted with ice-cold denaturing solution to a concentration of 0.2 mg/μL, followed by homogenization with BioMasher II (NIPPI, Japan) on ice. The lysate was disrupted using a Bioruptor sonicator (SONIC BIO Co., Japan) for 30 min (30 s on/30 s off, high setting) while being incubated in ice water, and then centrifuged the sample at 19,000 × g and 4°C for 30 min. The supernatant was collected, flash-frozen using liquid nitrogen, and stored at -80 °C. Mouse EDTA plasma was collected using MiniCollect^®^ □and immediately centrifuged at 1000 × g for 15 min at 4°C. Plasma samples were stored in aliquots at -80°C.

### Peptide extraction from mouse organs using modified DS method

Peptide extracts from mouse organs were prepared according to a previously described protocol [18]. Briefly, 50 μL of mouse organs supernatant was added dropwise to 1.5 mL of -80°C acetone (ACE). After centrifugation at 19,000 × g and 4°C for 30 min, the pellet was resuspended in 85% acetonitrile (ACN) in 12 mM HCl, mixed for 10 min at 4°C, and then sonicated for 20 min. The mixture was centrifuged at 19,000 × g for 15 min at 4°C, and 900 µL of the supernatant was collected and lyophilized. The lyophilized sample was resuspended in 75 µL of methanol and mixed for 5 min at 4°C. Subsequently, 1,425 µL of ice-cold ACE was added and mixed for 10 min at 4°C. After centrifugation at 19,000 × g and 4°C for 30 min, the supernatant was discarded, and the pellet was lyophilized. The dried sample was redissolved in 100 µL of 0.02% lauryl maltose neopentyl glycol (LMNG) in 0.1% trifluoroacetic acid (TFA) [19], desalted using MonoSpin C18 column, and lyophilized. The lyophilized sample was stored at -80°C until LC-MS/MS analysis.

### Peptide extraction from mouse plasma using DS method

Mouse plasma for peptide extraction was prepared as previously described [8], with slight modification. First, 50 µL of plasma was lyophilized. The dried plasma was then resuspended in 100 µL of a denaturation buffer (7 M urea, 2 M thiourea, 50 mM DTT) and mixed for 10 min at 4°C. The sample was sonicated for 10 min. Subsequently, the denatured plasma was added dropwise into 1.8 mL of -80°C ACE, and the mixture was incubated for 1 h at 4°C with mixing. After incubation, the mixture was centrifuged at 19,000 × g for 15 min at 4°C. The supernatant was then removed, and 1 mL of 85% ACN in 12 mM HCl was added to the pellet. The sample was sonicated for 30 min and mixed for 1.5 h at 4°C to extract peptide components. Afterward, the mixture was centrifuged at 19,000 × g for 15 min at 4°C, and 900 µL of the supernatant was lyophilized. The sample was redissolved in 100 µL of 0.02% LMNG in 0.1% TFA and desalted using a MonoSpin C18 column. It was then lyophilized and stored at −80 °C until LC–MS/MS analysis.

### LC-MS/MS analysis

The desalted samples were redissolved in a solution of 0.02% dodecyl maltose neopentyl glycol in 0.1% TFA. Peptide extracts from organs were prepared to a final concentration equivalent of 100 µg of each organs/µL, and peptide extracts from plasma were prepared equivalent of 1 µL of plasma/µL. For analysis, 4 µL of organ sample and 6 µL of plasma samp was injected onto an analytical column, 75 µm × 25 cm nano HPLC capillary column (IonOpticks, Collingwood, Australia), heated to 60°C. Peptides were separated using a NanoElute 2 system (Bruker Daltonics, Bremen, Germany). The mobile phases consisted of solvent A (0.1% formic acid (FA) in water) and solvent B (0.1% FA in 100% ACN). The 60 min gradient consisted of 0 min 5% B, 2 min 7% B, 53 min of 31%B and 55 min 50% B at a flow rate of 300 nL/min and 56 min 70% B and 60 min 70% B at a flow rate of 400 nL/min. Eluted peptides were analyzed on a timsTOF HT (Bruker Daltonics) via Captive Spray II (Bruker Daltonics)

timsTOF HT parameters were set as follows: Capillary voltage, 1,600 V; MS1 and MS2 range, 100 - 1700; 1/*K0* range, 0.75 – 1.25; dry gas, 3.0 L/min; and dry temperature, 180 °C. The collision energy was set by linear interpolation between 59 eV at an inverse reduced mobility of 1.60 versus/cm^2^ and 20 eV at 0.6 versus/cm^2^. To calibrate ion mobility dimensions, two ions from the Agilent ESI-Low Tuning Mix were selected (m/z [Th], 1/K0 [versus/cm2]: 622.0289, 0.9848; 922.0097, 1.1895).

For ddaPASEF mode, the parameters were as follows: Ramp and accumulation time; 100 ms, PASEF Number; 10, Charge range, 2-5; Target Intensity, 20,000; Intensity Threshold, 1,000. To achieve a 300 Hz sequencing speed, the measuring time was set to 2 ms and the switching time to 1.2 ms. Singly charged precursor ions were excluded using a polygon filter.

For diaPASEF mode, a modified Thin-diaPASEF was used [20]. The isolation window scheme was focused on divalent precursor ions in the m/z range of 350 to 1,100. This scheme was defined by 31 windows, each with an isolation width of 25 Th and an overlap of 1 Th between adjacent windows. The ramp and accumulation time was set to 75 ms, resulting in a total cycle time of approximately 1.0 s. The denoising mode was set to ‘Low Sample Amount (Sensitive)’ in diaPASEF data reduction.

### Empirical spectral library generation

DDA-MS data were analyzed using FragPipe v23.0 (https://fragpipe.nesvilab.org/)[21], [22] with a nonspecific peptidome workflow against a mouse FASTA file containing 43,510 entries (UP000000589 + decoys, downloaded, 2025, April). MSFragger[23] source parameters were as follows: precursor mass tolerance, 15 ppm; Fragment mass tolerance, 0.02 Da; cleaveage, NONSPECIFIC; Peptide length, 7 – 50; Peptide mass range, 500 – 12,000; Max variable mods on a peptide, 3; variable modifications, Oxidation(M), Acetylation(N-term), Amidation (C-term), pyro-Glu from Q, pyro-Glu from E. For validation, MSBooster[24] in combination with Percolater was enabled and results were filtered applying a peptide FDR of 1%. An empirical spectral library was built using the Spectral library generation module with standard parameters[25]. The empirical spectral library obtained by analyzing plasma in injection triplicate was designated as “Plasma”, while the empirical spectral library obtained by analyzing plasma in injection triplicate and eleven types of organs were designated as “Plasma and Organs”.

DIA-MS data were analyzed using DIA-NN v 2.3.1 (https://github.com/vdemichev/DiaNN)[26] with an InfinDIA pre-search against mouse FASTA file containing 43,510 entries (UP000000589, downloaded, 2025, April). The parameters for generating the empirical library were as follows: Protease, Non-specific; missed cleavage, 0; Max variable modifications, 3; peptide length range, 7–50; precursor charge range, 2–5; precursor m/z range, 350–1,100; and fragment ion m/z range, 100–1,700. ‘Prediction from FASTA’; ‘N-term M excision’; ‘ Acetyl (N-term)’ and ‘ Ox (M)’; were enabled and additional options; ‘--var-mod UniMod:2, -0.9840, *c’; ‘ --var-mod UniMod:27, -18.0106, E *n’ and --var-mod UniMod:28, -17.0265, Q *n’. The DIA-NN search parameters were as follows: MS1 accuracy, 15 ppm; MS2 accuracy, 15 ppm; protein inference and gene expression; Scoring, Proteoforms; Machine learning, NNs (cross-validated); Quantification strategy, Legacy (direct); Cross-run normalization, Off. The MBR was turned On. Results were filtered applying a precursor FDR of 1%. The spectral library obtained by analyzing plasma in triplicate was designated as “InfinDIA”.

### Peptide identification and quantification

DDA-MS data from plasma obtained in triplicate were analyzed using FragPipe v23.0 with a nonspecific peptidome workflow against a mouse FASTA file containing 43,510 entries (UP000000589 + decoys, downloaded, 2025, April). MSFragger parameters were the same as for empirical spectral library generation. MS1 quantification was performed using standard parameters by IonQuant.

DIA-MS data obtained from plasma in triplicate were analyzed using DIA-NN v 2.3.1 with library search mode using three types of empirical spectral library (Plasma, Plamsa and Organs, InfinDIA). Each analysis workflow was designated as DIA-Plasma (DIA-P), DIA-Plasma and Organs (DIA-PO), and DIA-InfinDIA (DIA-I). DIA-NN parameters were the almost same as for empirical spectral library generation, but the MBR was turned off.

### Statistical analysis and visualization

All statistical analyses were performed in R (version 4.5.2; R Foundation for Statistical Computing, Vienna, Austria). Data visualization was carried out using the ggplot2 package with R Studio.

## Results and Discussion

### Evaluation of analysis workflows for plasma peptidomics

Peptides were extracted from mouse plasma using the DS method, and a sample equivalent to 6µL plasma was analyzed with a 60-min gradient by DIA-MS and DDA-MS, each in triplicate. DIA-MS data were processed with DIA-NN v2.3.1 using three workflows (Fig. 1) described in Materials and Methods section (DIA-PO, DIA-P, and DIA-I), while DDA-MS data were processed with FragPipe v23.0 using the nonspecific peptidome workflow.

**Figure 1.**
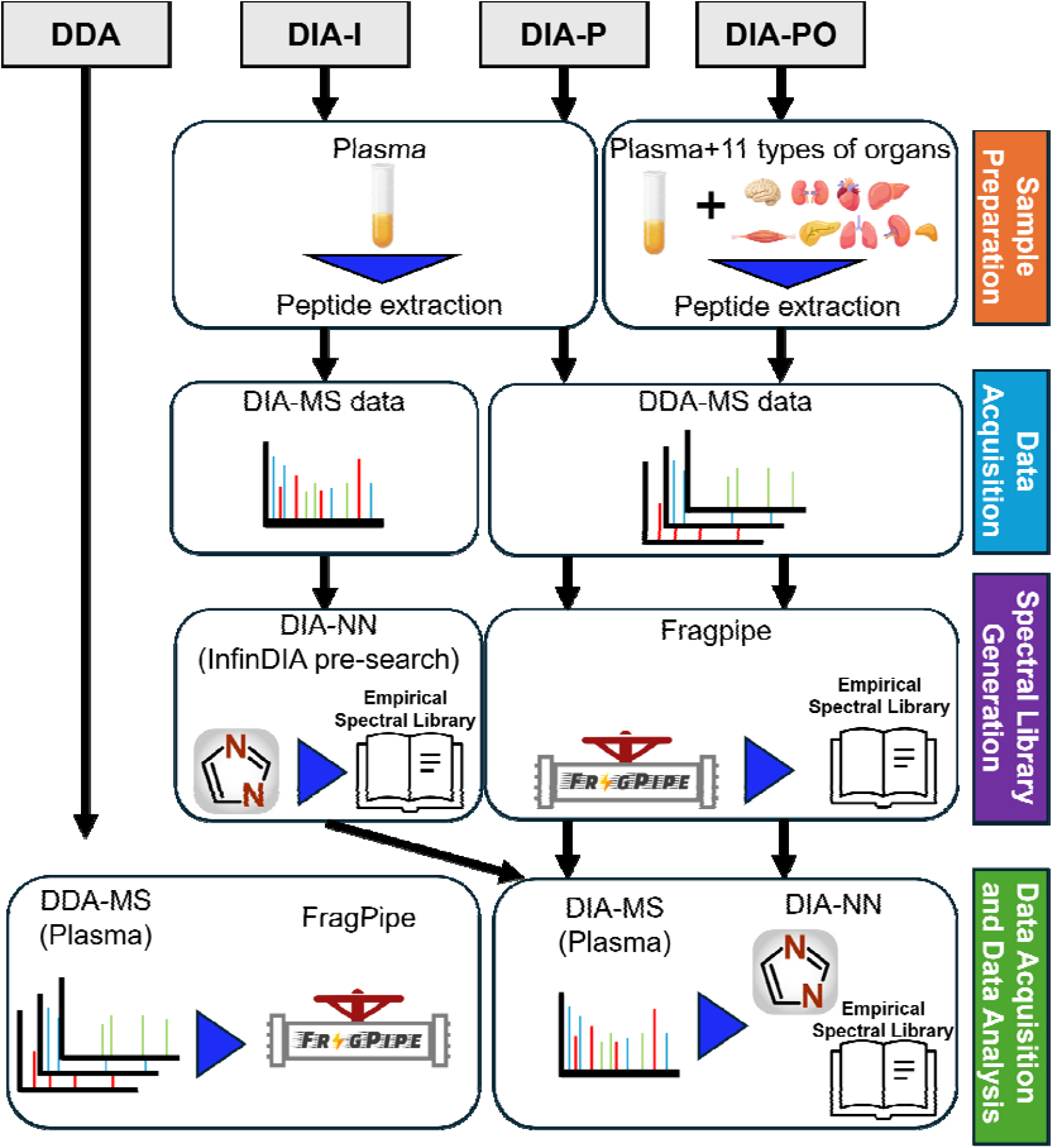
Workflow of DDA-MS analysis and DIA-MS analysis. DIA-I, DIA-P, and DIA-PO indicate DIA data analyzed using InfinDIA, the empirical spectral library of plasma, and that of plasma and organ, respectively.

Among the DIA analyses, only the DIA-PO workflow yielded a higher number of peptide identifications than DDA, identifying approximately 1.3-fold more peptides (5,587 peptides for DIA-PO vs. 4,394 for DDA) (Fig. 2A). Focusing on organ-derived peptides, which we defined as those identified in each of the eleven organs, DIA-PO detected approximately 1.6-fold more organ-derived peptides in plasma than DDA (2,108 vs. 1,287). Moreover, the number of peptides quantified across all three technical replicates was higher for DIA-PO than for DDA (4,717 peptides vs. 2,524 peptides, respectively). For organ-derived peptides specifically, 1,563 peptides were quantified with DIA-PO and 810 peptides with DDA (Fig. 2B). In the previous study by Lin Lin et al. [16], which represents the sole study combining plasma peptidomics with DIA-MS, an average of 2,302 peptides were identified, and 2,239 peptides were consistently identified across all three replicate measurements. In contrast, the DIA-PO workflow developed in the present study substantially outperformed these results, both in terms of the average number of identified peptides and the number of peptides identified in all three measurements. These findings suggest that the construction of an empirical spectral library from organ-derived peptides, together with peptide separation by trapped ion mobility spectrometry, has improved both detection sensitivity and technical reproducibility in plasma peptidomics.

**Figure 2.**
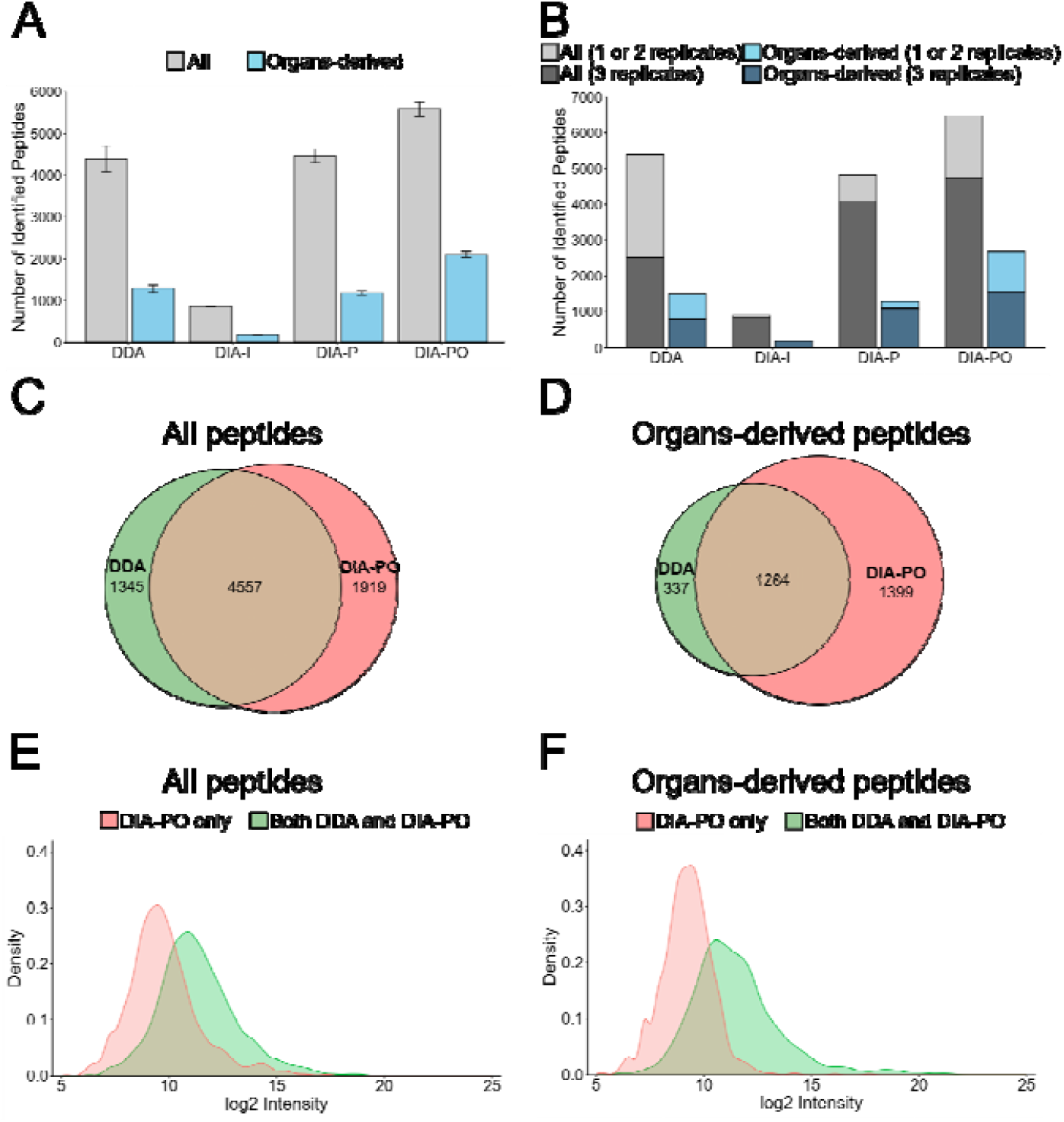
Analysis of peptides derived from an equivalent of 6 µL of mouse plasma extracted by the DS method. (A) The bar graph shows the average number of peptides identified across three technical replicates, with error bars indicating the standard deviation. “Overall” refers to all identified peptides. “Organ-derived” refers to peptides that were identified in at least one of the eleven organs. (B) Comparison of the reproducibility of peptide identification between analytical methods. Peptides were classified as quantified in all three technical replicates (dark gray or dark blue) or in one or two replicates (light gray or light blue). (C, D) Venn diagrams comparing peptides identified by DDA and DIA-PO. The numbers indicate peptides detected at least once across the three replicates. (C) All peptides. (D) Organ-derived peptides. (E, F) Distributions of log2-transformed peptide intensities for peptides detected at least once across three replicates, comparing peptides identified only by DIA-PO (red) with those commonly identified by both DIA-PO and DDA (green). (E) All peptides. (F) Organ-derived peptides.

The number of amino acid residues of the identified peptides tended to be slightly smaller in DIA-PO than in DDA, indicating that DIA-PO preferentially identified shorter peptides (Fig. S1). The distribution of charge among the identified peptides differed between DDA and DIA-PO, with doubly charged precursor ions being enriched in DIA-PO (Fig. S2). This trend reflects the measurement characteristics of Thin-diaPASEF [20], which is optimized for proteomics-style acquisition. We anticipate that further optimization of DIA parameters for peptidomics is expected to improve coverage beyond what was achieved in this study. The number of peptides identified in plasma was higher with DIA-PO than with DDA (Fig. 2C). Moreover, DIA-PO identified more organ-derived peptides in plasma than DDA (Fig. 2D). This observation suggests that incorporating an organ-derived empirical spectral library enabled more extensive identification of organ-derived peptides in plasma and, in turn, allowed us to better exploit the intrinsic high sensitivity of DIA, which had not been fully realized previously. DIA-PO-unique peptides tended to exhibit lower signal intensities than peptides detected by DDA-MS (Fig. 2E), and this tendency was even more pronounced for those that were also observed in organs (Fig. 2F). In addition, because organ-derived peptides, as expected, predominantly exhibited a low-intensity distribution. These results indicate that using an empirical spectral library of organ-derived peptides enables the detection of a larger number of peptides in plasma, particularly low-intensity organ-derived peptides.

As described above, the newly developed DIA-PO method improved the analytical sensitivity and quantitation of organ-derived peptides in plasma. However, the organ-derived empirical spectral library used in this study was generated from only eleven organs, each measured once by DDA-MS without reduction and alkylation of cysteine thiol groups (e.g., carbamidomethylation). In the future, expanding the spectral library by performing replicate measurements with reduction and alkylation and by including a broader set (or variety) of organs is expected to further increase the number of organ-derived peptides that can be comparatively analyzed in plasma.

### Organ-derived peptides identified in plasma

Using the DIA-PO workflow, we mapped 2,683 organ-derived peptides identified in plasma to their organs of origin (Fig. 3a). As a result, most peptides were observed in only a single organ, suggesting that many peptides present in the circulation may reflect the state of individual organs (Fig S3). Thus, by using this approach, it may be possible to determine which organs are associated with peptides whose levels change in plasma under various disease conditions. Among the identified bioactive peptides is neuropeptide Y (NPY), whose plasma concentration has been reported to be in the sub-pmol/L range [27]. To our knowledge, this is the first report of full-length NPY being identified in plasma by non-targeted analysis, supporting the high sensitivity of our approach.

**Figure 3.**
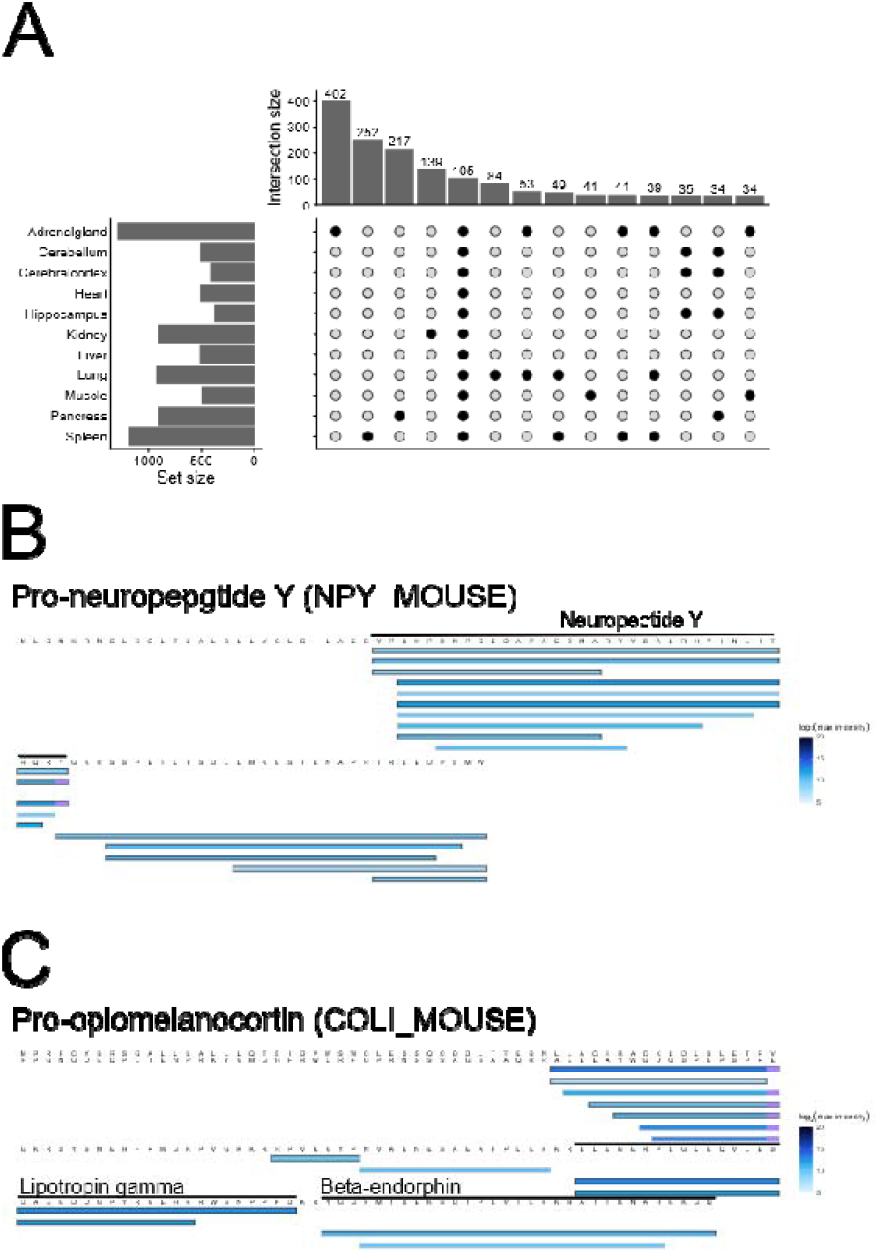
Contents of Organ-Derived Peptides in Plasma. (A) The UpSet plot shows the organ origins of organ-derived peptides detected in plasma. The figure shows only those with intersection size of 30 peptides or more. (B, C) Peptide alignment maps for prohormone precursor protein (B) NPY_MOUSE (UniProtID, P57774; Gene Name, Npy) and (C) COLI_MOUSE (UniProt ID, P01193; Gene Name, Pomc. The light blue squares indicate sites undergoing amidation. Color intensity reflects the maximum detection intensity for each peptide. Additionally, bars enclosed by black lines indicate organ-derived peptides, while those enclosed by gray lines indicate peptides not detected in organs.

The dataset also included 19 known bioactive peptides annotated as “Peptide” in UniProt (Table S1). These comprise established hormones and neurotransmitters, many of which were detected in secretory organs such as brain and pancreas.

Among all identified bioactive peptides, fragmented peptides lacking a few amino acid residues at either the N-or C-terminus were frequently observed (Figures 3B and 3C). A similar trend is known for peptides measured in organs [18]. Furthermore, among the non-organ-derived peptides identified by DIA-PO, 599 peptides were cleavage peptides of organ-derived peptides only detected by plasma DDA-MS data. Taken together, these findings suggest that secreted peptides undergo further processing or degradation in the circulatory system.

By combining DIA-MS with an empirical spectral library of organ-derived peptides, this workflow enables the distinction between peptides secreted from organs and those further fragmented in the blood. This feature should aid in the discovery of bioactive peptides and biomarkers, as well as in obtaining precise information about organs.

## Conclusion

This study proposed a novel strategy for detecting organ-derived peptides in plasma based on an empirical spectral library constructed from organ peptide analysis. As a result, comparative analysis of approximately 5,000 plasma peptides was achieved within about one hour of analysis. Approximately 2,000 of these peptides were organ-derived, and one-third of these were low-abundance peptides difficult to detect using conventional DDA methods or DIA methods employing empirical plasma peptide libraries. Among these were 19 known bioactive peptides, including neuropeptide Y at blood concentrations of several pM.

A key feature of this strategy is its ability to analyze plasma peptides associated with secretory organs. This enables the identification of causative organs for systemic diseases through plasma peptide analysis. Furthermore, it allows differentiation between peptides cleaved within the organ and those cleaved in the bloodstream after secretion. This is critically important for obtaining accurate information about secretory organs. The concept proposed in this study not only enables the discovery of novel bioactive peptides and biomarkers but also facilitates the monitoring of organ and tissue states through plasma analysis. The strategy presented here can be applied to humans through such as cultured cells and iPS cells, and is expected to significantly contribute to future life sciences and disease research.

## Author Contributions

Y.O., and Yoshio K. designed the study. M.I., T.T., and Takeshi M provided mouses samples. Y.O. and A.T. prepared peptidomic samples. R.K, Takashi M., and Yusuke K. equipped with computer systems and software. Y.O., R.K., and Yusuke K. performed LC-MS/MS measurements and analyzed the data. Y.O., Yusuke. K., and Yoshio K. wrote the paper. O.O., Yusuke K., and Yoshio K. supervised the project. All authors reviewed the results and approved the final version of the manuscript.

## Acknowledgments

This work was supported by JSPS KAKENHI Grant Number 17H02206 and 17K19926 to Yoshio K, JST, NBDC Grant Number JPMJND2304 to Yoshio K, and All Kitasato Project Study (AKPS) to Yoshio K, Takashi M, and Takeshi M.

## Data Availability

Mass spectrometry peptidomic data were deposited in the ProteomeXchange Consortium via the jPOST partner repository [28], with dataset identifiers PXD073758 for ProteomeXchange and JPST003835 for jPOST.

## Supporting Information

Additional supporting information can be found online in theSupporting Information section. SupportingInformationfile1:pmicxxxxx-sup 0001-SuppMat.docx SupportingInformationfile2: pmicxxxxx-sup-0002-Tables.xlsx

