## supplemental data for "DIA-MS based plasma peptidomic workflow for profiling organ-derived peptides"

**Contents :**


- Figure S1. Distribution of number of amino acid residues of peptides detected by DIA-PO and DDA.
- Figure S2. Charge-state distribution of precursor ions identified by DDA and DIA-PO.
- Figure S3. Distribution of organ-derived peptides identified in mouse plasma by DIA-PO according to the number of organs in which they were detected.

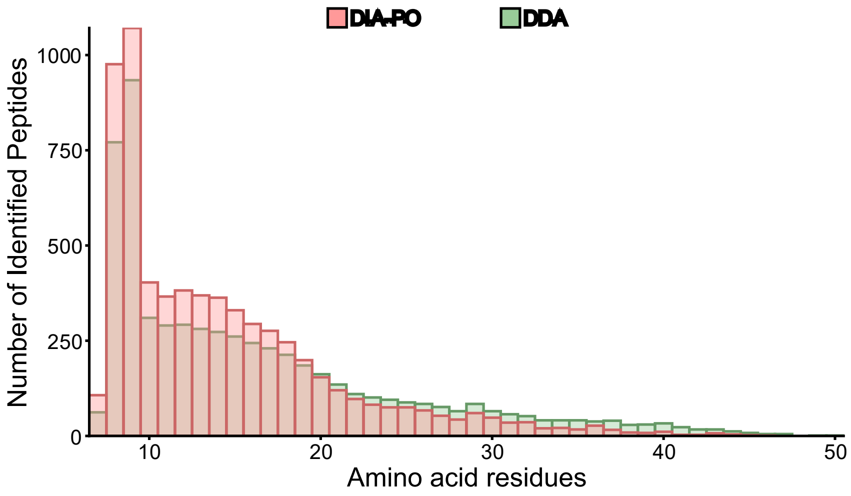


**Figure S1.** Distribution of number of amino acid residues of peptides detected by DIA-PO and DDA. All peptides identified across triplicate measurements for each method are shown (6,476 peptides for DIA-PO and 5,902 peptides for DDA).

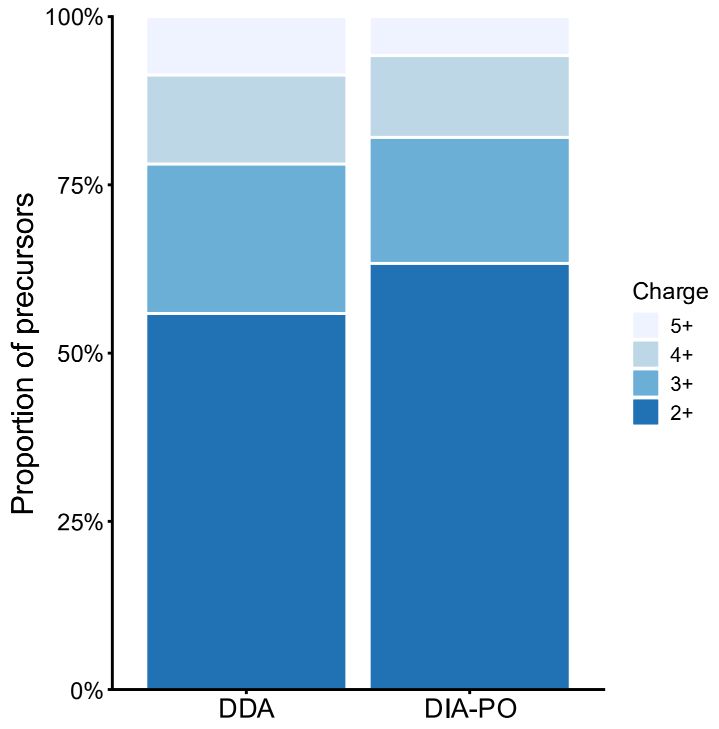


**Figure S2.** Charge-state distribution of precursor ions identified by DDA and DIA-PO. Stacked bars indicate the proportion of precursors with charge states 2+, 3+, 4+, and 5+ among all precursors identified across triplicate measurements for each method.

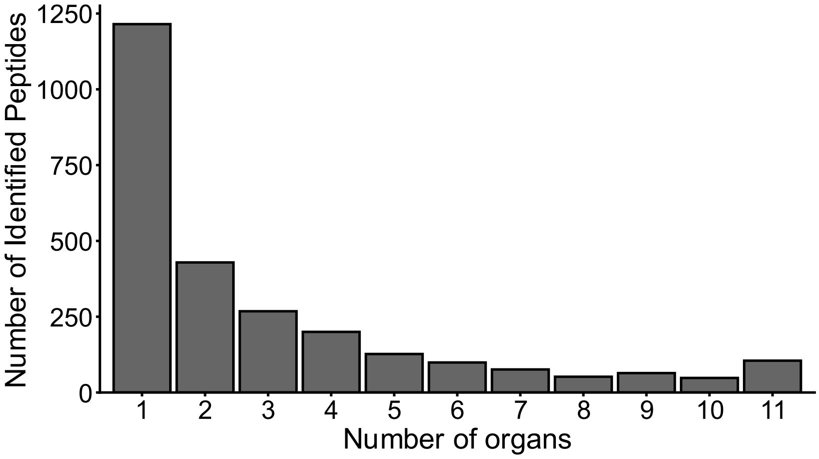


**Figure S3**. Distribution of organ-derived peptides identified in mouse plasma by DIA-PO according to the number of organs in which they were detected. The x-axis indicates the number of organs (1–11) in which a given peptide was identified, and the y-axis shows the corresponding number of peptides.
